# Dissecting Alzheimer’s disease heterogeneity by cross-trait polygenic prediction

**DOI:** 10.64898/2026.05.15.725551

**Authors:** William F. Li, Nabil Mohammed, David A. Bennett, Manolis Kellis, Yosuke Tanigawa

## Abstract

Mapping the genetic basis of inter-individual heterogeneity in multifactorial diseases opens the door to mechanistic insights and opportunities for targeted intervention. In Alzheimer’s disease (AD), clinical and pathological heterogeneity is well recognized, but genetic dissection is limited by a lack of well-powered cohorts with deep phenotypic characterization. Here, we introduce a polygenic score (PGS) analysis strategy to address these limitations by leveraging the inherent pleiotropy in complex trait genetics. We perform a cross-cohort, cross-trait application of pre-trained PGS, integrating 713 UK Biobank-derived PGS with 36 deep AD phenotypes across 1678 ROSMAP participants. We identify 268 statistically significant (FDR<0.1) associations between 12 prioritized PGS and 36 AD phenotypes. Prioritized PGS include blood lipid measurements, inflammatory biomarkers, and cancer traits; observed AD phenotypes include cognition, amyloid, and tangles. Of the 268 associations, 49 persist with *APOE*-excluded PGS. Predictive models trained on multiple prioritized PGS outperform the AD PGS or *APOE* alone for predicting amyloid and cognition. Lastly, our approach identifies six individual-level AD polygenic subtypes supported by distinct pathological patterns. Overall, we combine large-scale biobank resources and deeply-phenotyped cohorts using PGS, reveal genetic features underlying AD heterogeneity, and provide a general model for stratifying heterogeneous disease-focused cohorts using genomics.

## Introduction

Inter-individual heterogeneity is a fundamental feature of complex traits, where individuals with the same diagnosis can differ profoundly in their clinical course and underlying etiology^1,2^. For instance, Alzheimer’s disease (AD), the most prevalent neurodegenerative disease worldwide^3^, presents with significant variability in neuropathological markers^4,5^, cognitive decline patterns^6–8^, and disease genetics^9,10^. Notably, neuropathological markers, including neuritic plaques, diffuse plaques, and neurofibrillary tangles, differ in their regional distribution^11–13^, molecular correlates^14,15^, and clinical implications^16,17^. Differences in AD neuropathology are associated with variable treatment responses to modern disease-modifying therapies^18–23^. Mapping the genetic basis of AD phenotypes provides a scalable way to understand disease heterogeneity, predict individual-level manifestations, and inform clinical strategies tailored to patient subgroups.

Two complementary approaches provide promising insights into the biological basis of AD heterogeneity but fall short in directly connecting genetics with phenotypic variation. On the one hand, focused disease-specific cohort studies, such as the Religious Orders Study and Rush Memory and Aging Project (ROSMAP), have extensively profiled genetic variation^24,25^, cell-type-resolved molecular signatures^26–33^, neuropathologic measurements^25,34,35^, and cognitive change^36^ in thousands of aging individuals. On the other hand, large-scale genome-wide association studies (GWAS) on hundreds of thousands of individuals have revealed at least 75 AD risk variants and implicated biological processes^10,37–39^. However, these two existing approaches have insufficient statistical power to directly map the genetic contributions to AD phenotypic heterogeneity. Due to the clinical and technical obstacles in acquiring and profiling human tissue at scale, even the largest focused cohorts, such as ROSMAP, have orders of magnitude fewer individuals compared to the hundreds of thousands used in modern GWAS meta-analysis^40–46^.

Large-scale biobanks, such as the UK Biobank (UKB)^47–51^, have accelerated the discovery of common variant associations across thousands of traits using GWAS^49,52,53^. Polygenic scores (PGS) aggregate these variant-trait associations from hundreds to thousands of variants into a single score to estimate the genetic liability of a single trait^54,55^. For AD, individuals in the highest decile of disease-specific PGS distributions have up to a 1.9-fold greater risk of disease compared to those in the lowest decile^10,56^. However, single-trait PGS models capture only a fraction of the polygenic signal relevant to a given trait and cannot predict deep disease phenotypes not directly assessed in biobank-scale studies.

The pervasive pleiotropy in complex trait genetics offers a unique opportunity to investigate the genetic basis of phenotypes not directly measured in biobank-scale studies. Indeed, 20-60% of genetic associations are estimated to be pleiotropic^57–61^. For disease case-control prediction, leveraging pleiotropy using cross-trait PGS has led to improved predictive performance beyond what any individual single-trait PGS achieves^62–67^. Despite these advancements, little is known about whether cross-trait PGS approaches can be used to characterize the heterogeneity of a single disease. Motivated by this gap, we hypothesize that comprehensive PGS libraries trained on health phenotypes in biobank populations can be applied cross-trait to understand the heterogeneity of disease-related deep phenotypes in smaller, richly phenotyped cohorts.

Here, we address prior limitations and present an application of cross-cohort, cross-trait PGS analysis to leverage pleiotropy for the study of AD heterogeneity. Our analysis bridges 713 UKB-derived PGS^68^ with 36 observed deep AD phenotypes across 1678 ROSMAP participants^69^ to uncover cross-trait associations for deep phenotypes spanning the AD phenome. We report that many of the cross-trait associations reflect polygenic influences beyond the *APOE* locus. The genomic loci underlying prioritized PGS models are enriched for involvement in trait-relevant biological processes that facilitate the interpretation of the association results. We further show that combining information across multiple PGS improves the prediction of AD phenotypes and identifies six genetically defined subtypes marked by differing pathological patterns. Overall, our work highlights the contributions of polygenic signals to inter-individual variation in AD and demonstrates an application of PGS for dissecting the genetic basis of phenotypic heterogeneity in disease.

## Methods

### Compliance with ethical regulations and informed consent

We analyzed individual-level genetic and phenotypic data from the Religious Orders Study/Rush Memory and Aging Project (ROSMAP)^69^, and we derived previously characterized polygenic score (PGS) models from prior work on the UK Biobank resource^50,51,68^. The ROSMAP study consists of two longitudinal cohort studies of aging and dementia with extensive clinical and pathological phenotypic measurements (n=4067). Genotyping information is now available for nearly all autopsied participants^70^; we used an early subset of 1681 genotyped individuals^24^. *APOE* was directly genotyped^71^. Participants enroll without known dementia. Informed and repository consents were obtained from all participants, as well as an Anatomic Gift Act. An Institutional Review Board at Rush Medical Center approved both the Religious Orders Study and the Rush Memory and Aging Project^69^.

This research has been conducted using the UK Biobank Resource under Application Number 21942, “Integrated models of complex traits in the UK Biobank” (https://www.ukbiobank.ac.uk/enable-your-research/approved-research/integrated-models-of-complex-traits-in-the-uk-biobank). All participants of the UK Biobank resource provided written informed consent. More information is available at https://www.ukbiobank.ac.uk/explore-your-participation/basis-of-your-participation/.

### ROSMAP phenotype selection and processing

We obtained 126 clinical and pathological ROSMAP phenotypic variables described in the Rush Alzheimer’s Disease Center Research Resource Sharing Hub^69^ (**Table S1**), now with a total of 4067 individuals profiled for these phenotypes with a complete clinical evaluation. We used the 85 quantitative phenotypes. We focused on a subset of the individuals (n=1678) with both genotype (n=1681) and phenotype (n=4067) data for all downstream analyses.

To focus on ROSMAP phenotypes relevant to AD, we evaluated associations with two AD diagnosis variables and kept 37 phenotypic variables with statistically significant associations. We associated each ROSMAP phenotype using logistic regression with a binary clinical diagnosis (diagnoses made by a neurologist at the time of death following review of select clinical data without post-mortem data^72^) and binary NIA Reagan score for pathologic AD^73^. For each association, the diagnosis served as the dependent variable, while each ROSMAP phenotype was the independent variable. If a ROSMAP phenotype had p<1 x 10^-8^ significance in logistic regression in both logistic regression models, we determined that it was an AD phenotype and used it for further analysis. The TDP-43 pathology phenotype was generated as previously described^74^, resulting in a final count of 36 distinct AD-related quantitative phenotypes. For visualization, we performed Z-score standardization for each phenotype, ignoring missing values (**Figure S1**). For downstream analysis, we performed mean imputation across all 1678 joint dataset ROSMAP individuals for each of the resulting 36 ROSMAP phenotypes.

To characterize the phenotypic heterogeneity, we performed a PCA on the 36 variables over the 1678 ROSMAP individuals; for the input data to the PCA, we centered and standardized all variables to have a mean of 0 and a standard deviation of 1 over the 1678 individuals. We then quantified the variance explained by each component and visualized latent components using biplot visualization^75,76^, where we used scatter plots to show the ROSMAP individuals (observations) projected onto the PCs with overlaid arrows to indicate the PC loadings of each phenotype (variables) to the respective PCs (**Figure S3**).

### ROSMAP genotype quality control, liftOver, and imputation

For the ROSMAP genotype data, we obtained raw, unimputed genotype data from the Synapse Alzheimer’s disease (AD) Knowledge portal (accession ID: syn17008939)^77^. The genotyping was performed in two batches with the hg18 reference, one at the Broad Institute using Affymetrix GeneChip 6.0 for 1126 individuals with 749k SNPs per individual, and the other at the Translational Genomics Institute (TGen) for 582 individuals with 653k SNPs per individual^24^, for 1708 total individuals with genotype data. We performed quality control, liftOver, and imputation to prepare the genetic data on the hg19 reference, which was used in the PGS models in the UKB^68^ (**Figure S2**).

First, we applied a missingness filter on the individuals with a 10% threshold, which filtered out 7 individuals, for a total of 1701 individuals (1120 from the Broad, 581 from TGen). We applied a missingness filter on the variants with a 10% threshold.

Second, we converted the genomic coordinates from hg18 to hg19 using the variant ID (rsID) in dbSNP^78^ (version 155) and aligned the alleles using the GRCh37 reference genome^79^. For multi-allelic sites, we kept the variant at the site only if one of the alleles matched the hg19 reference sequence. We confirmed that 99.5% of the variants in the original hg18 dataset were mapped to the hg19 coordinates. We additionally checked our genotyping data against the Haplotype Reference Consortium, which showed 99.7% rsID match^80,81^.

Third, we applied genotype imputation followed by an additional round of quality control. To that end, we split the dataset into two batches based on the SNP Array batch each individual belonged to (Broad or TGen). Within each batch, we imputed the genotype data through the Michigan Imputation Server with Eagle 2.4.1 phasing and Minimac4 imputation^82–84^. The imputation pipeline defines about 160 sub-chromosomal chunks (161 for Broad and 157 for TGen) and checks the chunk-level missingness of the input genotype for each individual. We removed a total of 20 individuals (14 from Broad and 6 from TGen), each of whom had more than 50% variant missingness in at least one of the 160 chunks. We performed genotype imputation for autosomes and chromosome X separately: the 21 individuals with over 50% missingness in a chromosome X chunk only (10 from Broad and 11 from TGen) have no imputed chromosome X genotype data, but are retained in the dataset; there is additionally 1 retained individual from the Broad batch with no imputed chromosome 20 genotype data. Overall, after imputation, we have a total of 1681 individuals, consisting of 1106 from the Broad batch with 39.6 million imputed variants and 575 from the TGen batch with 38.2 million imputed variants. Of the 1681 individuals, 21 have no chromosome X genotype data, and 1 has no chromosome 20 genotype data.

To inspect the genetic ancestry of individuals, we used pre-computed principal component analysis (PCA) loadings of genetic variants from the Human Genome Diversity Project and the 1000 Genomes Project as a reference^85–87^ and projected ROSMAP individuals passing genetic quality control into the latent principal components (PCs) (**Figure S4**).

### Polygenic scoring of ROSMAP individuals

We used sparse UKB PGS models from our previous study, one of the largest collections of PGS models trained on the same cohort with the unified PGS training procedure^68^. In the previous study, we applied the Batch Screening Iterative Lasso algorithm^88^ to 269,704 unrelated white British individuals in the UKB resource, resulting in PGS models for a range of traits, including disease outcomes, cancer registry data, family history for disease, blood biomarkers, and imaging-derived measures, such as volumes of the gray matter in specific brain regions^50,68^. We analyzed PGS models reported with statistically significant predictive performance (nominal p<2.5 x 10^-5^), except for those for time-to-event traits, resulting in 713 PGS models analyzed in the study (**Table S2**). Among the two PGS models trained for AD-related phenotypes, we focused on the PGS model trained for family history of AD (n_case_=41,451 of 269,704 in UKB) as the primary PGS model for AD (“AD PGS”), given its superior statistical significance in the UKB hold-out test set (p=9.8 x 10^-107^ vs p=6.3 x 10^-28^) and broader genetic basis (139 underlying genetic variants vs 15 underlying variants). In contrast, “prostate cancer PGS” throughout this paper refers primarily to the PGS model trained for prostate cancer risk alone rather than the one for family history, given the superior statistical performance in this case without family history (p=8.9 x 10^-87^ vs p=2.8 x 10^-40^).

We computed PGS for each of the 1678 ROSMAP individuals using each of the 713 PGS models in our UKB-derived PGS library (**Figure S5a**). We performed the computation using PLINK v2.00^89^ to find the average score over the alleles, with missing variants imputed by the mean allele frequency.

### Transferability analysis of UKB AD PGS for prediction within ROSMAP

To assess the transferability of UKB PGS models on ROSMAP, we assigned each of the ROSMAP individuals a binary label of AD (NIA Reagan score<=2) or non-AD (NIA Reagan score>2). We excluded individuals with missing NIA Reagan scores from this analysis (407 individuals excluded, with 1271 remaining for the analysis). We computed the receiver operating characteristic (ROC) curve for AD PGS in predicting the binary AD label as well as the area under the ROC curve (AUROC) (**Figure S6**).

### Covariates used for cross-trait association analysis

To adjust for covariates in the two-stage cross-trait association, we included age at baseline, age at death, sex, a binary variable for the batch in which genotyping was performed (TGen or Broad), and the 10 leading population PCs (**Table S1**). We computed population PCs using PLINK v2.00^89^, the Human Genome Diversity Project and the 1000 Genomes Project reference weights^85,86^, and publicly available scoring scripts^87^.

### Two-stage cross-trait polygenic association

We conducted cross-trait analysis using UKB-derived PGS and observed ROSMAP AD phenotypes using a two-stage procedure. First, to select PGS relevant to AD, we performed the first stage of cross-trait association. We selected five global observed AD phenotypes (amyloid density, tangle density, global pathology, global cognition level, and global cognition resilience slope). For each trait (both UKB PGS and observed in ROSMAP), we performed z-score standardization across the ROSMAP individuals. We then performed a pairwise linear regression between each of the 5 observed phenotypes as the dependent variable and each PGS in our entire library of 713 PGS as the independent variable, with the described covariates. For each pairwise linear regression, we recorded the effect size (standardized regression coefficient) of the association, the standard error in the effect size estimate, and the p-value of the linear relationship (**Figure S7, Table S3**). We computed the false discovery rate (FDR) over the p-values of all 3565 (= 5 ROSMAP phenotypes * 713 UKB PGS) pairwise associations using the Benjamini-Hochberg procedure^90^. We retained PGS with at least one pairwise association at FDR<0.5, yielding a collection of 12 AD-relevant PGS (**Figure 2a**).

**Figure 1.**
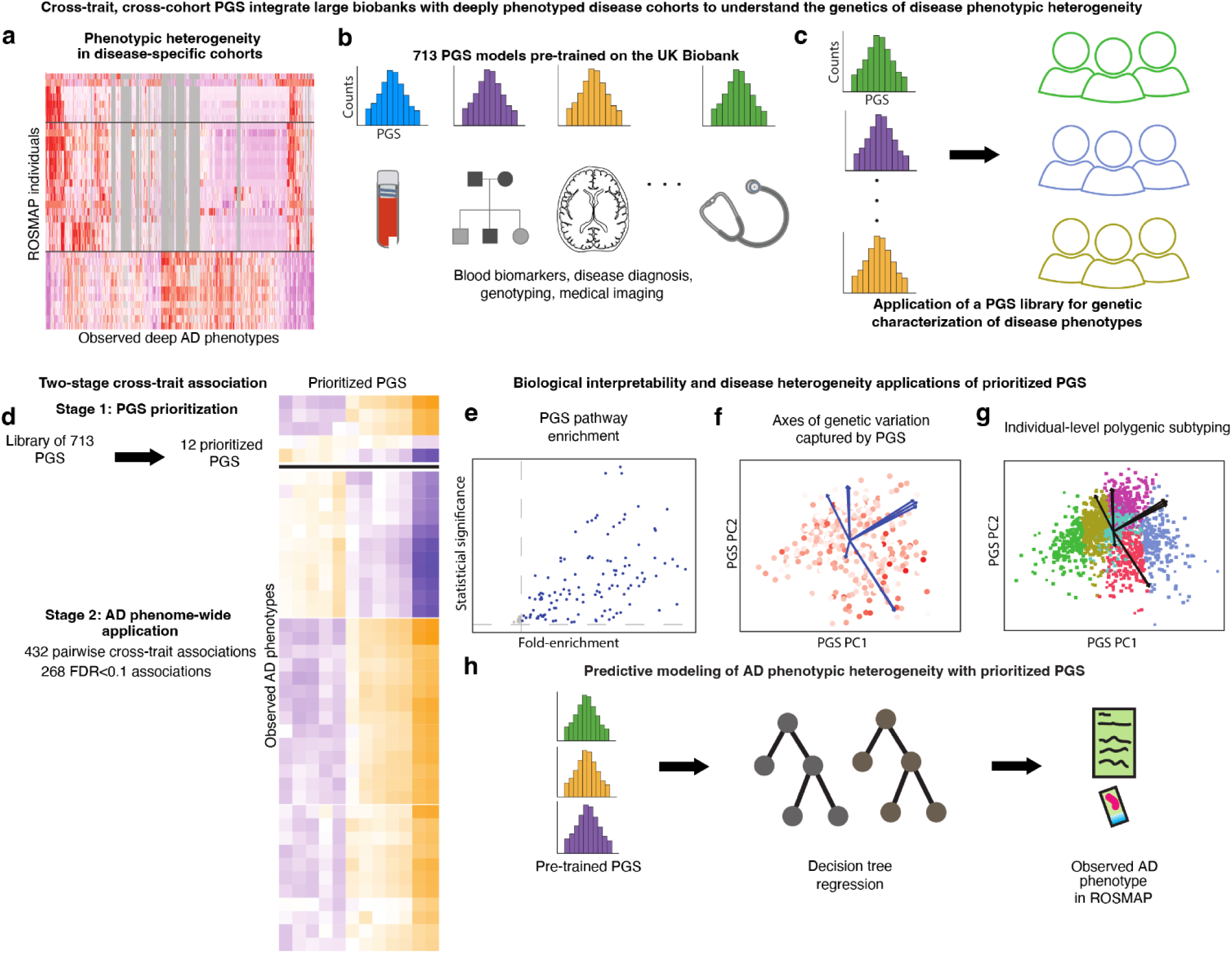
Overview of the study. **(a)** Alzheimer’s disease presents with cognitive and histopathological phenotypic heterogeneity. **(b)** We used polygenic score (PGS) models across 713 traits characterized from the UKB resource. **(c)** A cross-trait application of a PGS library pre-trained on the UKB enables characterization of the genetics of AD phenotypic heterogeneity across individuals. **(d)** Using computed PGS and observed ROSMAP cohort phenotypes, we performed a two-stage phenome-wide association analysis to characterize the genetic basis of phenotypic heterogeneity in AD. In Stage 1, we used the cross-trait association to prioritize relevant PGS as genetic features. In Stage 2, we use prioritized PGS to characterize observed AD phenotypes in ROSMAP. **(e)** We further characterized prioritized PGS via GREAT enrichment on leading PGS model loci to identify enriched biological pathways. **(f)** Principal components (PCs) were computed for prioritized PGS, and loadings of each PGS were used to visualize the distinct axes of prioritized PGS. **(g)** We performed individual-level genetic subtyping of the ROSMAP cohort using PGS PCs. **(h)** We performed predictive modeling to evaluate the predictive power of multiple-PGS genetic features for the prediction of observed AD phenotypes.

**Figure 2.**
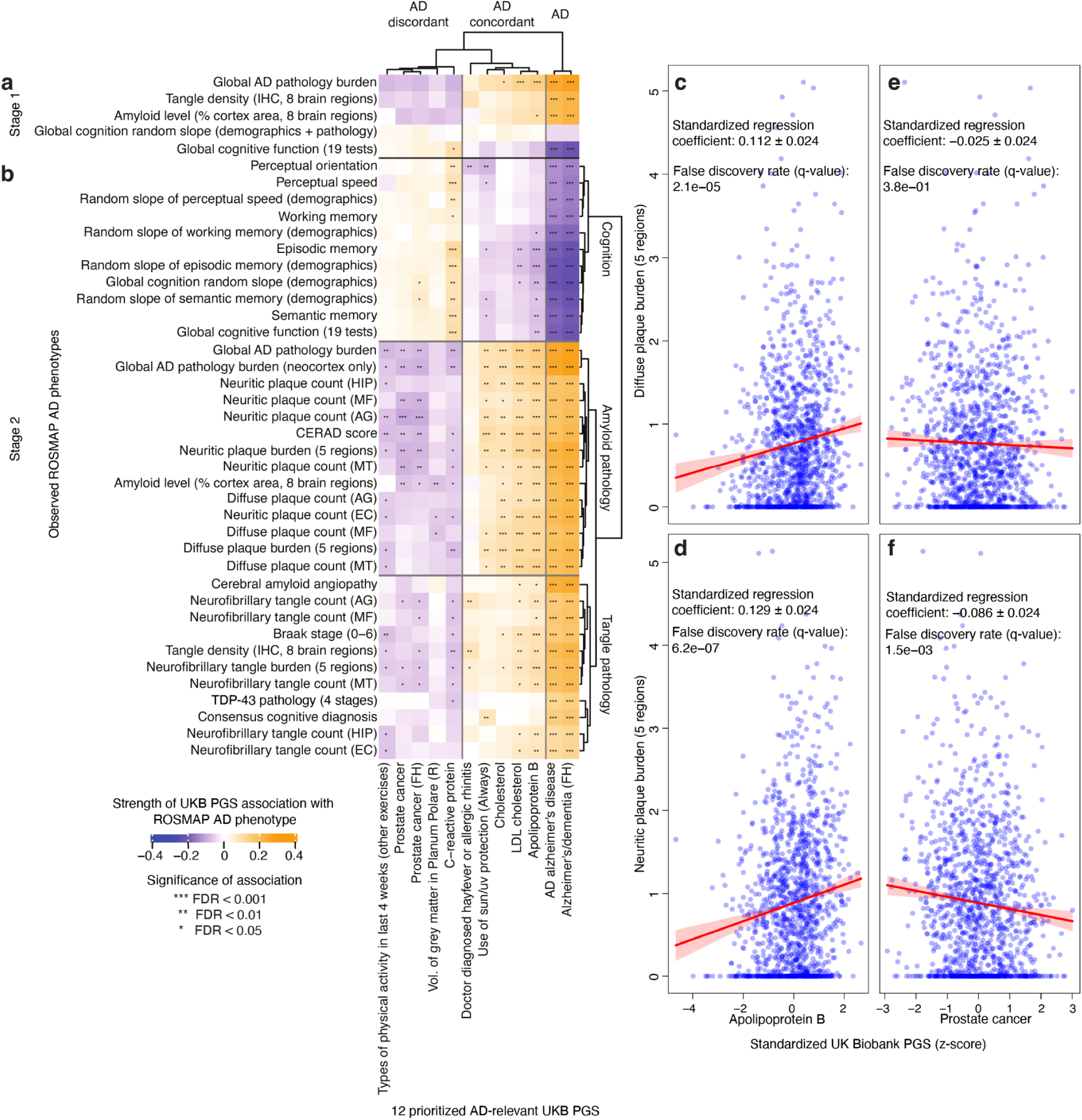
Cross-cohort cross-trait application of UKB PGS to ROSMAP AD phenome reveals genetic basis for phenotypic heterogeneity in AD. **(a-b)** We show associations of the 12 most relevant PGS (x-axis) with 5 global **(a)** and 36 distinct **(b)** observed AD phenotypes (y-axis), each sorted by hierarchical clustering (dendrograms). The color represents the strength of association between pairs of PGS and observed AD phenotypes, with asterisks marking statistically significant associations. **(c-f)** We show ROSMAP individuals plotted by PGS z-scores (x-axis) and observed plaque burden (y-axis). We show the line of association through the centroid of the data points, where the slope of the line is given by the effect size computed in the stage 2 analysis, and the shaded error regions are 95% confidence intervals (**Methods**).

To thoroughly characterize AD-related phenotypes, we performed the second stage of cross-trait association. We calculated pairwise associations between the 12 PGS and 36 observed AD phenotypes using multivariable linear regression, with the covariates as described above. For each variable (both UKB PGS and observed ROSMAP phenotype), we performed z-score standardization across the ROSMAP individuals. For each pairwise linear regression, we recorded the effect size of the association, the standard error in the effect size calculation, and the p-value of the linear relationship (**Figure 2b, Table S4**). We computed the FDR over the p-values of 432 pairwise associations (= 36 ROSMAP phenotypes * 12 UKB PGS) using the Benjamini-Hochberg procedure.

To visualize specific pairwise associations from the second stage of cross-trait association, we plotted individuals by observed ROSMAP phenotype value and standardized z-score PGS (**Figure 2c-f**). We furthermore plotted a line through the centroid of the individual-level coordinates with slope calculated from the pairwise association effect size; in particular, because the phenotype values are not z-score normalized in this plot, the slope shown in the plot is the recorded effect size (i.e., standardized regression coefficient) from **Table S4** multiplied by the standard deviation of the observed ROSMAP phenotype over the individuals. The shaded region around the plotted line was computed as a 95% confidence interval by multiplying the standard error from **Table S4** by 1.96, then likewise multiplying by the standard deviation of the observed phenotype.

### *APOE-*exclusion association analysis

To evaluate the impact of the *APOE* region on our findings, we excluded the genetic variants in the *APOE* locus (chr19:45,176,340–45,447,221, hg19) from each UKB PGS model, following the literature^91–94^, and repeated the second stage of cross-trait association using the modified 12 PGS, recording effect sizes, errors, p-values, and FDR calculated with Benjamini-Hochberg over the 432 p-values (**Figure 3a,c,e,g,i, Table S5**).

**Figure 3.**
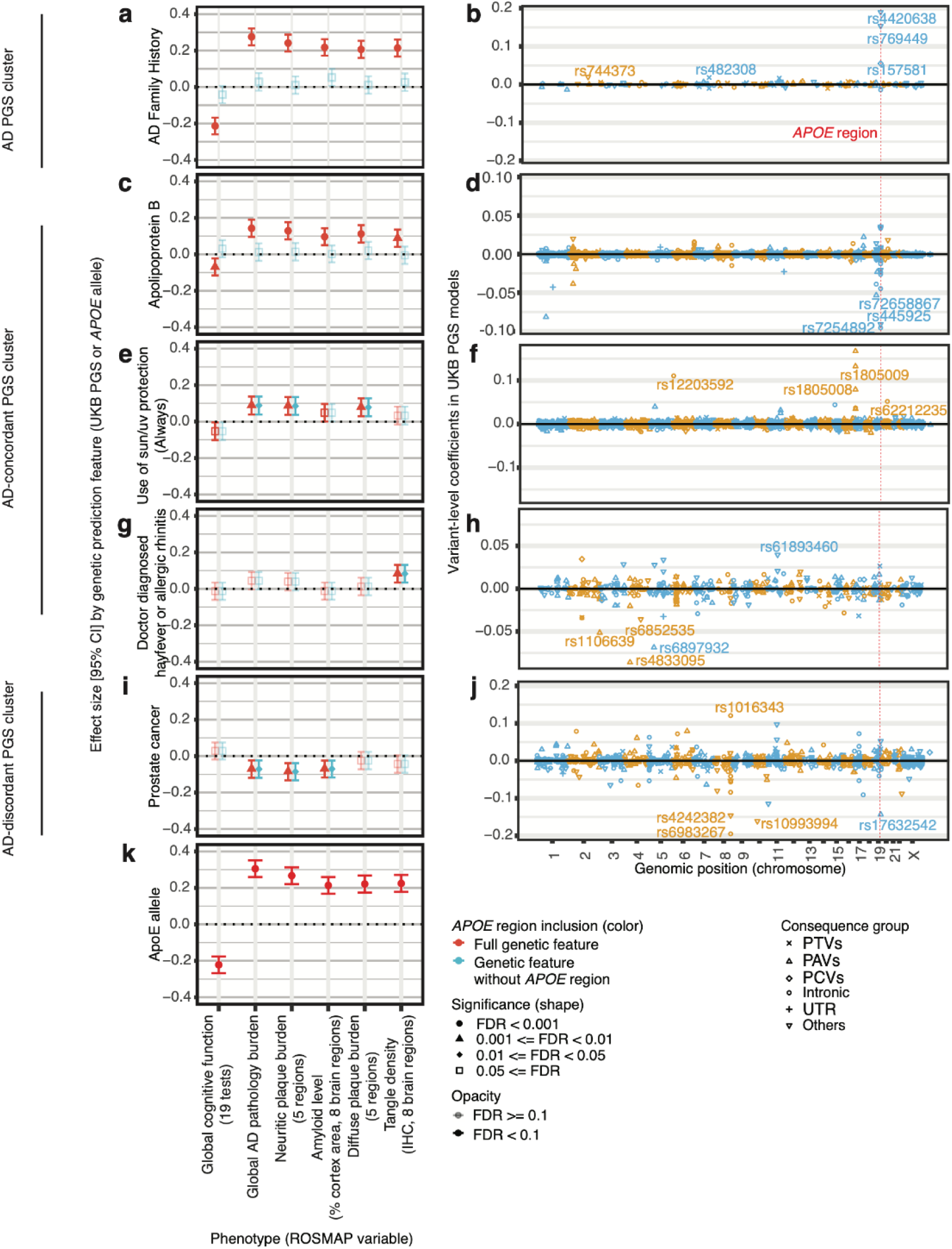
The effect of the *APOE* allele on cross-trait associations between PGS and observed AD phenotypes. **(left)** We show association profiles for individual PGS, showing the effect sizes (y-axis) between each genetic feature (a PGS or *APOE*) and a selection of 6 observed AD phenotypes (x-axis). Error bars show the 95% confidence interval. Each plotted effect size is colored by whether its respective genetic feature has an *APOE* contribution, and with shapes and opacity based on statistical significance. **(right)** We show effect sizes by model weights (y-axis) over the span of the genome (x-axis) for the respective PGS models. We annotate representative variants with large absolute effect sizes by rsID. Vertical red dotted lines demarcate the *APOE* region. Shapes of points in the plots are consequence groups. PTV, protein-truncating variant; PAV, protein-altering variant; PCV, proximal coding variant; UTR, untranslated region. Shown are PGS for AD **(a)**, ApoB **(c)**, use of sun/UV protection **(e)**, hayfever or allergic rhinitis risk **(g)**, prostate cancer **(i)**, as well as the *APOE* allele **(k)**. We also show the PGS model coefficients (y-axis) by genomic position (x-axis) for the prioritized PGS, with each variant annotated by consequence group (color). We show the UKB PGS models for AD **(b)**, Apolipoprotein B **(d)**, Use of sun/UV protection (always) **(f),** Doctor diagnosed hayfever or allergic rhinitis **(h)**, and Prostate cancer **(j).**

We additionally encoded *APOE* as a continuous *APOE*-gradient variable using the following mapping for defining a gradient *APOE* variable (*APOE*-gradient) ^95^: allele ε2/ε2 maps to -2, ε2/ε3 maps to -1, ε2/ε4 maps to -0.5, ε3/ε3 maps to 0, ε3/ε4 maps to 1, and ε4/ε4 maps to 2. Of the 1678 ROSMAP individuals we are analyzing, we found 5 individuals have missing *APOE* genotype values; we imputed these 5 using the mode genotype (ε3/ε3). We performed an association between *APOE*-gradient and the 36 AD-related observed ROSMAP phenotypes using multivariable linear regression with the described covariates. We recorded the effect sizes, errors, p-values, and Benjamini-Hochberg FDR over the 36 p-values (**Figure 3k, Table S6**).

### Biological characterization of PGS variants

To better understand the AD-relevant PGS, we downloaded the UKB PGS model weights^68^ for prioritized PGS, and we plotted variant coefficients in these models by genomic position (**Figure 3b,d,f,h,j**). We queried the implicated biological functions of top loci using the Open Targets Platform^96^.

For each of the prioritized AD-relevant PGS, we took the 1000 loci in the respective PGS model with the greatest absolute weight or all the loci if there are fewer than 1000 as the representative loci, and subsequently applied the genomic regions enrichment of annotations tool (GREAT, v4.0.4)^97,98^ to evaluate the statistical enrichment of biological processes in Gene Ontology (GO)^99,100^. We filtered the GO biological processes by hypergeometric FDR<0.05, then displayed them by binomial FDR and region-fold enrichment (**Figure 4a-b, Table S7-S8**).

**Figure 4.**
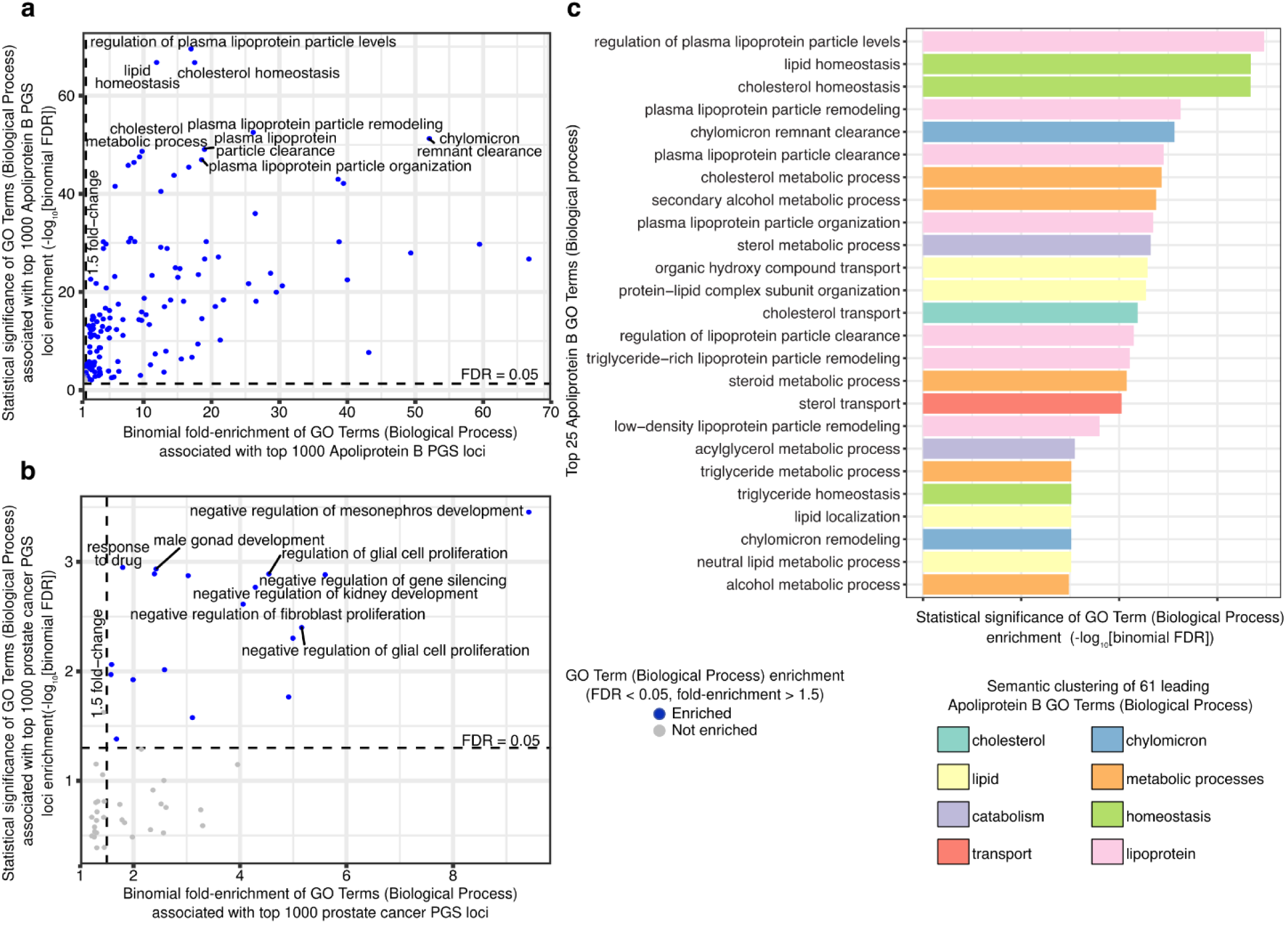
Genomic loci in prioritized PGS models are enriched for relevant biological processes in Gene Ontology (GO). To evaluate the biological relevance of prioritized PGS models, we applied the Genomic Regions Enrichment of Annotations Tool (GREAT) on the top 1,000 loci from two example PGS models (**Methods**). (**a**-**b**) We show the binomial fold enrichment (x-axis) and the statistical significance (binomial FDR) (y-axis) for the PGS models for ApoB (**a**) and prostate cancer (**b**). We highlight the significant (FDR < 5%, horizontal dashed line) enrichment with >1.5 binomial fold-changes (vertical dashed line) in color. **(c)** Semantic clustering (color) of the top 25 most significantly (x-axis) enriched GO terms (y-axis) for the ApoB PGS model reveals eight key processes (**Methods**, **Figure S8**).

To group similar enriched GO terms, we performed semantic clustering using the Global Vectors for Word Representation (GloVe), a 50-dimensional word vector mapping pretrained on Wikipedia 2024 + Gigaword 5^101–103^. Specifically, we focused on enriched GO terms (hypergeometric FDR<0.05 and binomial FDR<10^-15^) and removed words with limited semantic meaning (e.g., “of”, “to”, “-”). To obtain the semantic vector of a GO term, we averaged word-level vector representations for each component word. We applied k-means clustering on the semantic representation of GO terms, resulting in clusters of semantically similar GO terms (**Figure 4c, Figure S8**). In our ApoB PGS model analysis, we manually identified a unifying biological theme for the eight identified clusters.

### Multiple-PGS predictive modeling and principal component genetic feature computation

To evaluate the potential of a multiple-PGS model in predicting observed ROSMAP AD phenotypes, we compared the performance of our selected PGS with known genetic features implicated in AD. We split the 1678 individuals randomly into training (70%, n=1174) and testing (30%, n=504) sets. To generate an additional set of genetic features and to better understand the space of PGS, we performed a PCA with centering and standardization of the 12 AD-relevant PGS for individuals in the training set to generate PC loadings (**Figure 5a-b**). We computed PGS principal components (PCs) among the test set by projecting test set individuals onto these PCs (**Figure S5b**). We performed a biplot visualization^75,76^, showing the PC scores of the test set individuals as a scatterplot overlaid with arrows showing the contributions of each PGS to the respective PC (**Figure 5c-e)**. We chose the specific PCs to show on the biplot first by variance explained (PC1, PC2) and then by identifying a PC that we noted to have a strong contribution from PGS with *APOE*-independent association findings (PC6).

**Figure 5.**
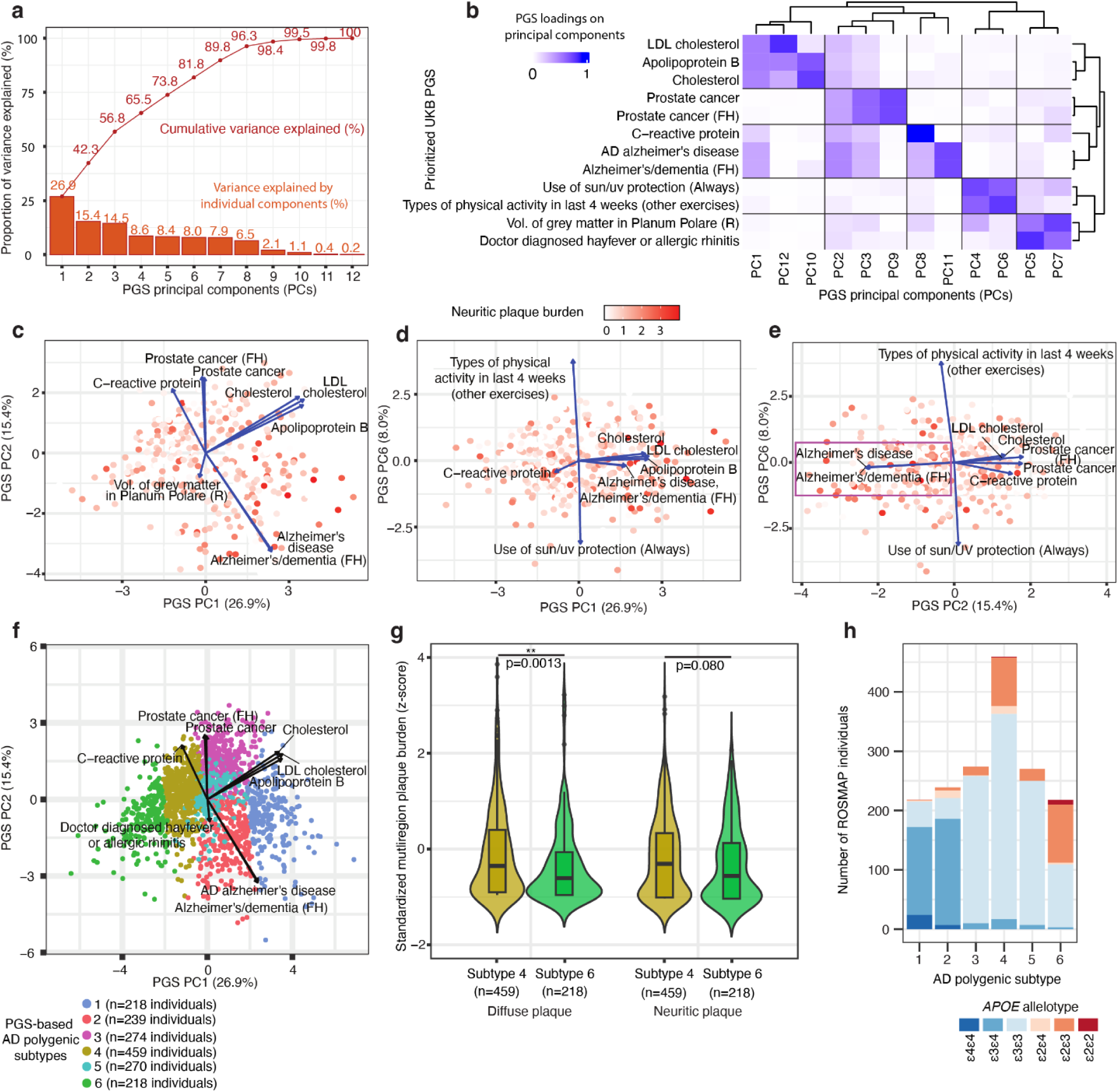
Prioritized PGS inform phenotypic heterogeneity in AD. **(a)** For principal components (PCs) characterized from 12 prioritized PGS (PGS PCs) (x-axis), we show the proportion of variance explained by each component (y-axis) in the training set (n=1174, 70%). We show individual components in the bar plot and cumulative variance explained in the line plot. **(b)** For each PGS PC (x-axis), we show the relative importance of contributing PGS (y-axis) as absolute PCA loadings (color). We show hierarchical clustering as dendrograms. (**c**-**e**) Biplot representing the projected PC scores of test set (n=504, 30%) individuals (dot) and the PGS loadings for each PC (arrows). The color represents the phenotypic value of neuritic plaque burden. We show PC1 vs. PC2 in (**c**), PC1 vs. PC6 in (**d**), and PC2 vs. PC6 in (**e**), selected based on the overall variance explained and the relevance of the *APOE* loci in the PC (**Methods**). Violet outline in (**e**) emphasizes that PC2 separates AD PGS from cholesterol PGS. **(f)** We show PGS-based AD polygenic subtype assignments of all 1678 ROSMAP individuals (color) as a biplot of PGS PC1 (x-axis) and PGS PC2 (y-axis), with overlaid PGS arrows as in **c**. **(g)** We show the distribution of observed AD phenotype value (color) within each AD polygenic subtype for selected variables (x-axis). We used the Wilcoxon rank-sum test to compare the distributions for each variable (**Methods**). **(h)** For each PGS-based AD polygenic subtype (x-axis), we show the number of ROSMAP individuals (y-axis) stratified by the *APOE* allelotype (color).

We then compared five different modeling schemes for the prediction of the variation of six observed AD phenotypes and two overall AD diagnosis criteria using extreme gradient boosting (XGBoost)^104^, training 40 total predictive models. In the first scheme, we trained the XGBoost predictors using only covariates. In the second, we used the quantified *APOE* allele. In the third, we trained the predictors using AD PGS only. In the fourth, we used all 12 AD-relevant PGS from the Stage 1 association. In the fifth, we used the top 8 PCs computed from PCA of PGS values in the training set. Each scheme included the previously described covariates as additional predictor features for the observed AD phenotypes. In each scheme, we trained 8 predictors, one for each global AD phenotype or AD diagnosis criterion, and used 200 rounds of boosting to train each predictor with default parameters otherwise.

To evaluate the predictors trained in each scheme, we first used each predictor to predict values for its global observed AD phenotype among test set individuals, and we then computed the Pearson correlation coefficient between predicted observed AD phenotype values and ground truth values (**Table 1, Table S9**). We computed Shapley values for the contribution of each genetic feature in the multiple-PGS prediction schemes (**Figure S9**).

**Table 1.**
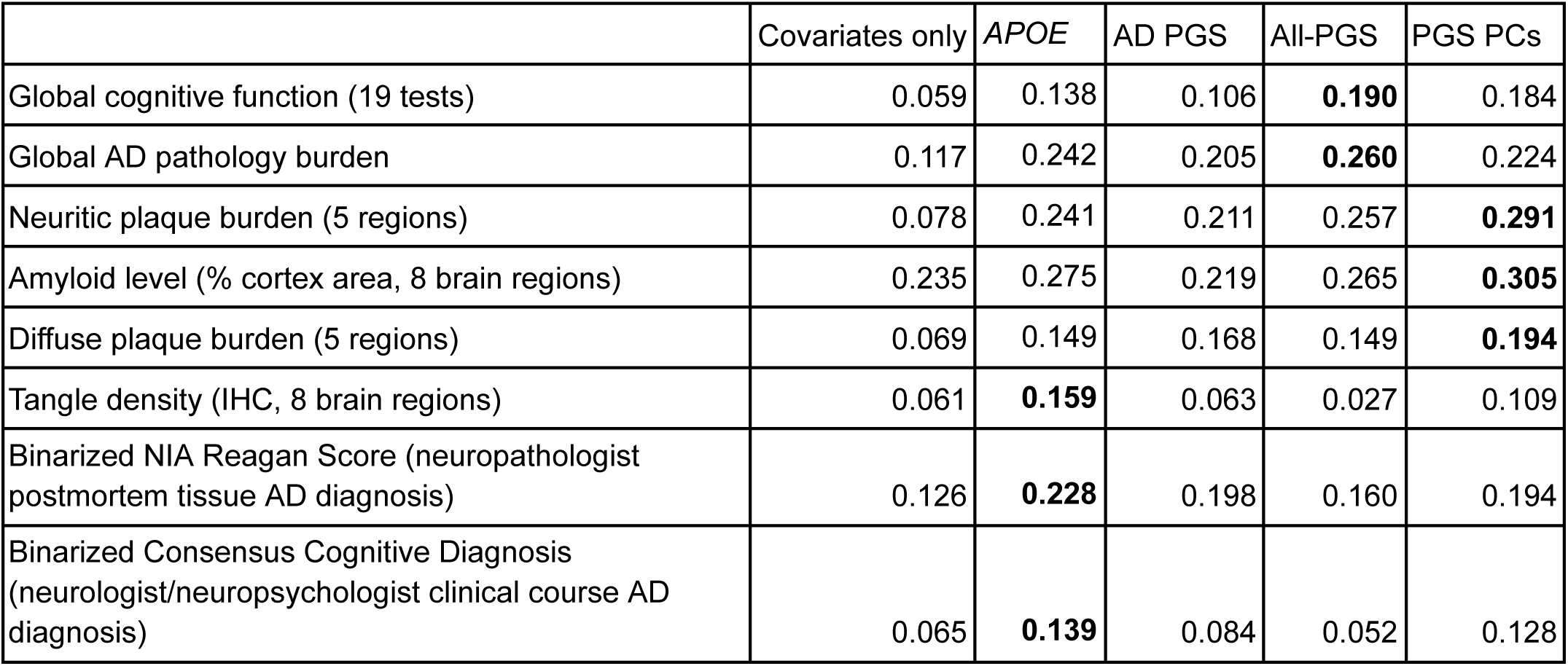
Predictive performance of gradient-boosted models. . We evaluated the predictive performance (Pearson’s correlation *r*) of gradient-boosted models with different predictor variables (columns) across six observed AD phenotypes and two compiled AD diagnosis variables (rows). *APOE* indicates a continuous apolipoprotein E genotype variable (*APOE* gradient), and PGS PCs indicate the top 8 principal components of prioritized PGS (**Methods**).

### PGS-based individual subtyping

We used genetic features derived from the 12 prioritized AD-relevant UKB PGS to propose genetic subtypes of AD liability. We generated PGS PCs weighted by relevance by multiplying each of the PGS PC values for each individual by the calculated variance-explained values (**Figure 5a**). Using these relevance-weighted PCs, we used *k*-means clustering to propose and assign individuals to six individual-level subtypes (**Figure 5f**), termed “AD polygenic subtypes”. For each subtype, we calculated the average subtype value for selected ROSMAP phenotypes (**Figure S10**). We performed a Wilcoxon rank-sum test^105^ on selected variable values between two chosen subtypes, indicating the computed p-values on the plot (**Figure 5g**). We additionally calculated the number of individuals with each *APOE* allelotype (**Figure 5h**). We stratified individuals by *APOE* allelotype and repeated the Wilcoxon rank-sum test between subtypes within selected allelotypes (**Figure S11**). In the calculation of average phenotype values, we used the z-score-standardized scaling as evaluated for (**Figure S1**) visualization.

## Results

### Study design to investigate the genetic basis of AD phenotypic heterogeneity

We analyzed 1678 individuals with genotyping and phenotyping of 36 distinct observed AD-related cognitive and histopathological phenotype variables (the “AD phenome”) in ROSMAP (**Figure S1-S3, Table S1**). Those individuals exhibited substantial phenotypic heterogeneity in cognition, amyloid, and tangles. Applying principal component analysis (PCA), we found that different AD phenotypes have distinct loadings along the top 2 principal components (PCs), with global amyloid variables having opposite and weaker contributions to PC1 compared to global cognitive function (**Figure S3b**). Overall, the leading twelve phenotypic PCs explained 40.1% of the overall variation (**Figure S3a**).

To complement detailed phenotypic characterization in ROSMAP with genetic predictors characterized from much larger cohorts, we turned to 713 PGS models pre-trained over 269,704 individuals in the UK Biobank^68^ (**Figure 1b, Figure S5a, Table S2**). Our approach leveraged both the depth of AD-specific phenotyping in ROSMAP and the statistical power of genetic information from the UKB by computing UKB PGS in ROSMAP individuals to generate biologically relevant genetic features (**Figure 1c**). To validate the application of UKB PGS in ROSMAP, we evaluated the cross-cohort predictive performance of the AD PGS based on family history (**Methods**). The AD PGS achieved an AUROC of 0.577 for observed NIA Reagan Score AD diagnosis in ROSMAP individuals (**Figure S6**), performing comparably with the AUROC of 0.558 in the UKB hold-out test set and confirming comparative predictive performance across cohorts.

With evidence of transferability, we designed a two-stage cross-trait association procedure to characterize the genetic basis of the AD phenome using UKB PGS models (**Figure 1d**). In Stage 1, we screened all 713 PGS against five global observed AD phenotypes. We used a permissive threshold (FDR<0.5) to minimize false negatives in candidate selection. In Stage 2, we used the prioritized PGS from Stage 1 to characterize the full set of 36 observed AD phenotypes with multiple-testing correction. We further performed biological pathway enrichment analysis on the top genomic loci in prioritized PGS models (**Figure 1e**), characterized the genetic variance captured by those PGS via PCA (**Figure 1f**), and applied them to genetic prediction and individual-level subtyping of the AD phenome (**Figure 1g-h**).

### Cross-trait UKB PGS associations to ROSMAP phenotypes reveal the genetic basis of phenotypic heterogeneity in AD

To assess the polygenic basis of AD-relevant multivariate phenotypes, we systematically assessed the 713 UKB PGS for their predictive performance for 36 observed phenotypes among 1678 ROSMAP individuals. Specifically, we applied a two-stage procedure to maximize the statistical power (**Methods**). Briefly, we selected five observed global AD phenotype variables, including amyloid, tangles, combined pathology, cognition, and slope of cognitive decline, representing the major axes of phenotypic variation in ROSMAP (**Figure S7**, **Table S3**). In the Stage 1 association analysis, our analysis prioritized 12 PGS with suggestive associations (FDR<0.5 with at least one of the 5 phenotypes) (**Figure 2a**). We used the permissive threshold to minimize type II error. In the Stage 2 analysis, we assessed the pleiotropic associations between prioritized PGS across a wider range of observed phenotypes; we expanded the set of observed phenotypes to the full set of 36 phenotypes. Overall, we found 268 statistically significant cross-trait associations (FDR<0.1) across 12 PGS and 36 observed phenotypes (**Figure 2b**). At more stringent FDR thresholds, we found 233 cross-trait pairs (FDR<0.05), 173 pairs (FDR<0.01), and 112 pairs (FDR<0.001). Hierarchical clustering of association summary statistics revealed three clusters among PGS (AD, AD concordant, and AD discordant) and three clusters among observed phenotypes (cognition, amyloid pathology, and tangle pathology).

Our analysis revealed differential patterns of association for PGS across the AD phenome, with each cluster of observed phenotypes having a unique set of enriched genetic features. We noted strong correlations of the AD PGS across the AD phenome, with FDR<0.001 associations for all of the 36 observed AD phenotypes. Among observed phenotypes in the cognition cluster, the PGS for C-reactive protein and Apolipoprotein B (ApoB) showed the strongest correlations, with 10 and 7 distinct pairwise FDR<0.05 associations with observed cognition variables, respectively. For observed phenotypes in the amyloid pathology cluster, the most strongly associated PGS were cholesterol, LDL cholesterol, and ApoB, which each had FDR<0.05 associations with 14 out of 14 of the observed variables in the cluster. Finally, for the tangle pathology cluster, the strongest associations were with the PGS for LDL cholesterol (8 out of 11 associations with FDR<0.05), ApoB (8 out of 11), and C-reactive protein (7 out of 11). Notably, cholesterol PGS had an FDR=0.0497 association with the observed 5-region neurofibrillary tangle burden and an FDR=0.0414 association with the observed Braak stage, the only associations it had with FDR<0.05 in the tangle pathology cluster. We also found a specific example of a revealed differential pattern of association. While ApoB PGS had FDR<0.01 associations with both observed diffuse plaque burden (**Figure 2c**, FDR=2×10^-5^) and observed neuritic plaque burden (**Figure 2d**, FDR=6×10^-7^), prostate cancer PGS had an FDR<0.01 association only with observed neuritic plaque burden (FDR=0.0015) and not with observed diffuse plaque burden (**Figure 2e-f**, FDR=0.375). This example contrasted a genetic correlate for both types of plaque burden together (ApoB PGS) with a genetic correlate for neuritic plaque burden alone (prostate cancer PGS).

### Distinguishing *APOE*-dependent and independent genetic correlations

To understand the effect of the *APOE* locus, we next compared cross-trait associations with and without *APOE*. Overall, we identified 49 cross-trait pairs that remained significantly associated (FDR<0.1) without the *APOE* locus. For example, associations that involved AD and ApoB PGS relied heavily on *APOE* contributions (**Figure 3a-d, Table S5**), whereas associations that involved sun/UV protection and prostate cancer PGS maintained associations independent of *APOE* (**Figure 3e-j, Table S5**).

Without *APOE*, only observed perceptual orientation retained an FDR<0.1 (FDR=0.07) association with AD PGS; by contrast, with the *APOE* region, AD PGS had FDR<0.001 associations across nearly the entire AD phenome. Similarly, without *APOE*, ApoB PGS had no FDR<0.1 associations for any observed AD phenotype. We additionally noted that the *APOE* allele itself shows an association pattern with the AD phenome resembling that of AD PGS risk with *APOE*, highlighting the strength of the *APOE* contribution to the AD PGS risk estimate (**Figure 3k, Table S6**).

Conversely, several other PGS identified in Stage 1 did not exhibit this dependence. Use of sun/UV protection and hayfever or allergic rhinitis PGS retained multiple FDR<0.1 associations over the AD phenome (**Table S5**). We noted that the use of sun/UV protection PGS retained FDR<0.05 associations across observed amyloid pathology variables (**Figure 3e**, e.g., FDR=0.02 for overall neuritic plaque burden and FDR=0.03 for overall diffuse plaque burden), and hayfever or allergic rhinitis PGS retained FDR<0.05 associations for tangle pathology variables (**Figure 3g**, e.g., FDR=0.02 for tangle density). Similarly, each of the PGS in the AD-discordant cluster, other than PGS for C-reactive protein, retained FDR<0.1 associations for 2 or more observed variables in the AD phenome (**Table S5**). Prostate cancer PGS retained its association strength across observed variables, particularly for neuritic plaque (**Figure 3i**, FDR=0.02 with overall neuritic plaque burden). Therefore, we found that both of the major pathological clusters (amyloid and tangles) of the observed AD phenome retained statistically significant cross-trait associations without the inclusion of *APOE*.

Because each PGS model linearly summed coefficients from individual variants, and we used sparse PGS models, we were able to interpret the coefficients for each PGS. Qualitatively, AD PGS (**Figure 3b**) and ApoB PGS (**Figure 3d**) were both enriched for loci in the *APOE* region, while prostate cancer PGS (**Figure 3j**), for instance, did not show similar dependence, consistent with our findings from our *APOE* exclusion analysis. At a more granular resolution, the strongest effect size outside of the *APOE* region for the prostate cancer PGS was from rs6983267^106–108^.

### Analysis of top variants for each PGS reveals enriched biological processes

We next asked whether aggregating the top genomic loci by effect size for each prioritized PGS could identify enriched biological pathways linking PGS to observed AD phenotypes. Using the genomic regions enrichment of annotations tool (GREAT)^97,98^, we found that the genomic regions comprising the ApoB PGS model were enriched for annotations related to lipid metabolism and lipoprotein regulation (**Figure 4a, Table S7**), processes that have been noted to be implicated in amyloid pathology^109–114^ and may provide a putative biological interpretation of the association results between ApoB PGS and observed amyloid pathology (**Figure 2b**). On the other hand, the genomic regions comprising the prostate cancer PGS model were enriched in annotations for kidney development, male gonad development, glial cell proliferation, and fibroblast proliferation (**Figure 4b, Table S8**).

To look into the composition of the Gene Ontology (GO) annotations enriched for the ApoB PGS model, we embedded each annotation by semantic meaning using a pre-trained word vector model (**Methods**), and we clustered GO terms using their semantic meaning vectors. We identified the clustered GO terms to fall under eight biological themes, and we show the 25 leading GO terms by their statistical significance of enrichment and biological theme (**Figure 4c, Figure S8**). We observed the four leading GO terms fall in the homeostasis and lipoprotein themes.

### Multiple-PGS models enhance the prediction of multiple AD-relevant phenotypes

Having prioritized PGS associated with multiple AD phenotypes, we next asked whether joint modeling of these PGS could improve the prediction of observed deep AD phenotypes, an approach informed by prior work on multiple-PGS-based prediction^62,63^. We trained nonlinear gradient-boosted models to predict six key AD phenotypes and two overall AD diagnosis variables. We evaluated four sets of genetic features alongside covariates, as well as a covariates-only baseline model. For the two benchmarks, we used the *APOE* allele and AD PGS. We then trained a model using all 12 prioritized PGS from Stage 1 (**Figure 2a**) as the genetic features. We also trained a model using the top 8 PGS PCs from a PCA of the training set (**Figure 5a**); test set PCs were computed by projecting onto the training set PCs. On the test set, the all-PGS model achieved the best predictive performance for global cognitive function and overall AD pathology, while the PGS-PCs model performed best for the three amyloid-related variables (**Table 1, Table S9**). *APOE* had the best performance for tangle density and overall AD diagnosis variables. Across AD phenotype prediction tasks, the multiple-PGS models relied on distinct sets of contributing PGS predictors (**Figure S9**), showing PC1 as the leading contributor for most phenotype predictions whereas PC2 was the leading contributor for consensus cognitive diagnosis and global cognitive function.

### Identified PGS characterize heterogeneity at the individual level in an aging and dementia cohort

Having validated the utility of cross-trait PGS application in recovering the genetic basis of deep AD phenotypes at the population level, we investigated prioritized PGS for characterizing both the phenotypic and genetic heterogeneity among ROSMAP individuals. To account for shared genetic effects across the prioritized PGS, we randomly split the n=1678 ROSMAP individuals into a training set (n=1174, 70%) and a held-out test set (n=504, 30%) and performed a PCA of the 12 selected PGS scores in the 1174 training set individuals (**Methods**). We found that the top 8 PCs combined explained 96% of the variance. Moreover, each PC explained at least 5% of the inter-individual variance of the PGS (**Figure 5a-b**), indicating there were multiple orthogonal directions of genetic variance captured by the PCs.

We next jointly visualized phenotypic and genetic heterogeneity among ROSMAP individuals using multiple PGS. We projected the remaining 504 held-out test set individuals using the PCA loadings (**Figure S5b, Methods**). We showed these individuals as points on biplots^75,76^ (**Figure 5c-e**). In these biplots, each point represents an individual, with position determined by projected genetic PC scores derived from multiple PGS and color indicating overall neuritic plaque burden.

The biplots recapitulated known relationships between neuritic plaque burden and AD genetic risk, with higher observed plaque burdens noted toward the right-hand side (higher PC1) of both plots (**Figure 5c–d**). PC1 had strong positive contributions from PGS for AD, cholesterol, LDL cholesterol, and ApoB and negative contributions from PGS for C-reactive protein and prostate cancer risk (**Figure 5c-d**). This visualization reflected our earlier quantitative finding that observed neuritic plaque burden correlates with the identified PGS (**Figure 2b**). On the other hand, for projections onto PC2 by PC6 (**Figure 5e**), we noted no such qualitative pattern of neuritic plaque burden; indeed, based on the PCA loadings arrows on the biplot and our quantitative results in (**Figure 2b**), we would not expect any.

We next used the prioritized PGS to propose individual-level polygenic subtypes of AD liability. Using the 12 computed PCs weighted by proportion of variance explained (**Methods**), we clustered the 1678 ROSMAP individuals to propose six PGS-based AD polygenic subtypes (**Figure 5f)**. We noted, as expected, that individuals within each of the six subtypes are qualitatively closely grouped when plotted on a PC1 vs PC2 biplot. We further showed the number of ROSMAP individuals assigned to each of the six subtypes, as well as the *APOE* allelotype compositions and selected observed phenotype averages for each subtype (**Figure 5g-h, S10**). Notably, we found that individuals within each subtype exhibit a diversity of *APOE* allelotypes, and hence, the *APOE* allelotype alone would not be sufficient to capture the heterogeneity proposed by these subtypes. While subtypes 4 and 6 differed in their observed diffuse plaque burden distributions with p<0.01 by the Wilcoxon rank-sum test, that statistically significant difference was not present for neuritic plaque burden (**Figure 5g**). On the other hand, with both observed diffuse and neuritic plaque burden, we found subtype 6 had a lower median than subtype 4, a trend maintained in three out of four comparisons after stratifying individuals within each subtype by *APOE* allelotype (**Figure S11**).

## Discussion

Here, we present a genetic dissection of phenotypic heterogeneity in complex traits using a systematic application of cross-trait polygenic score (PGS) analysis, using Alzheimer’s disease (AD) as a case study. We combine 713 PGS models pre-trained on 269,704 UKB individuals and 36 deep AD phenotypes across 1678 ROSMAP individuals. We report 268 statistically significant associations between PGS and observed AD phenotypes (FDR<0.1). Gradient-boosted models integrating multiple PGS demonstrate improved predictive performance for amyloid and cognitive AD phenotypes relative to baseline genetic predictors. Finally, we group individuals into AD polygenic subtypes and observe subtype-specific differences in observed diffuse plaque pathology.

In the context of human genetics, our study illustrates how predictive modeling with pleiotropy can be used to map the genetic basis of phenotypic heterogeneity within disease. Previous studies have employed pleiotropy to enhance the predictive performance of PGS in case-control disease risk prediction^62,63^. Here, we introduce an application for mapping the genetic basis of disease heterogeneity, leveraged through a large collection of biobank-trained PGS^68^. The proposed cross-cohort, cross-trait strategy is particularly attractive when a densely phenotyped disease-focused cohort has a smaller sample size compared to the cohorts used in conventional case-control GWAS and meta-analyses, as is the case with ROSMAP. We envision similar approaches to help characterize the genetic basis of phenotypic heterogeneity in other multifactorial diseases^115,116^.

Our cross-trait analyses demonstrated two specific promising use cases for applying PGS resources from large-scale cohorts to disease-focused cohorts with deep phenotypic characterizations: improved genetic prediction of disease phenotypic heterogeneity and a genetics-based subtyping of individuals. First, combining multiple AD-relevant PGS into a joint gradient-boosted predictive model outperformed single-score approaches for amyloid and cognition phenotypes, suggesting a generalizable way to more accurately predict deep AD phenotypes from genetics alone, consistent with prior findings^62,63^. Second, the same set of PGS was used to cluster individuals into distinct genetic subtypes, which exhibit both phenotypic differentiation and diverse *APOE* allelotype compositions. Together, these results suggest that integrating cross-trait polygenic signals may provide a useful framework for refining genetic risk stratification and for exploring genetically defined subtypes of AD liability in future studies.

Biologically, our study identifies genetic factors associated with distinct aspects of the AD phenome. We identify genetic features beyond *APOE* and AD PGS: of the 268 significant PGS-phenotype associations, 49 (18.3%) remained significant after excluding the *APOE* locus entirely. Notably, significant associations with both observed amyloid and tangle pathologies persisted, indicating that non-*APOE* genetic variation informs both major neuropathological axes of AD. We reveal three main clusters of AD phenotypes arising from association patterns: cognition, amyloid pathology, and tangle pathology variables. Likewise, we reveal three main clusters of AD-relevant PGS: PGS that directly predict AD (“AD PGS”), PGS whose association patterns are concordant with those of AD PGS, and PGS whose association patterns are discordant with those of AD PGS. Significant associations emerge between PGS and AD phenotypes for every combination of PGS cluster and phenotype cluster, showing a diversity of genetic factors associated with each dimension of the AD phenome.

Several PGS-AD phenotype associations warrant follow-up analysis on their causal relevance. For example, we found prostate cancer PGS retained associations specific to observed neuritic, but not diffuse, plaque burden, suggesting genetic dimensions of heterogeneity between neuritic plaque formation and diffuse plaques. GREAT^97,98^ enrichment of the prostate cancer PGS model loci showed pathways involved in glial cell and fibroblast proliferation regulation, suggesting these gene sets as directions for follow-up to evaluate their potential role in neuritic plaque formation. We also found that the hayfever or allergic rhinitis PGS retained FDR<0.05 associations specifically with observed tangle pathology variables, generating biological hypotheses for follow-up on whether neuroinflammatory pathways differentially modulate tangle burden^117–119^.

We note several directions for future study. First, because each PGS model represents a weighted combination of genetic variant effects, cross-trait associations can highlight biological pathways for mechanistic follow-up, for example, GREAT-enriched lipid metabolism pathways in relation to neuritic plaque burden. Second, our results rely on genetic correlations between PGS and observed AD phenotypes and do not establish causality. The association between ApoB PGS and observed amyloid pathology, for example, does not demonstrate that LDL-related variants causally affect amyloid burden, as shared genetic architecture could reflect horizontal pleiotropy or correlated downstream effects. Causal inference through pleiotropy-aware Mendelian randomization or related methods will be needed to evaluate specific mechanistic hypotheses arising from these associations^120,121^. Third, the PGS models used here were trained in European-ancestry individuals from the UKB^68^, and ROSMAP is itself predominantly of European ancestry^69^. Further studies are needed to generalize our findings to distinct genetic ancestry groups. Finally, further replication, preferably in prospective cohorts, is the next step to validate and refine our multiple-PGS predictive models and genetic subtypes.

Overall, our results establish an application of cross-trait PGS for pairing population-based genetic resources with deeply phenotyped disease cohorts to characterize disease heterogeneity. Our strategy addresses the statistical power limitations arising from the limited sample sizes of richly phenotyped disease cohorts. In our application to AD, the systematic mapping of *APOE*-dependent and -independent genetic contributions across amyloid pathology, tau pathology, and cognition domains provides a foundation for future mechanistic investigations of disease heterogeneity. The multiple-PGS predictive framework and individual-level genetic subtyping presented in our work have potential for developing genetic risk stratification efforts with larger and more diverse cohorts and deeper integration with molecular and functional data.

## Acknowledgments

We acknowledge general support from National Institutes of Health (NIH) grants AG054012, AG058002, MH109978, AG062377, AG081017, NS129032, AG077227, NS110453, NS115064, AG062335, AG074003, NS127187, AG067151, MH119509, HG008155, DA053631, R01-AT011460, and R56AG081376 (to M.K.). ROSMAP is supported by P30AG10161, P30AG72975, R01AG17917, R01AG015819, U01AG072572, and U01AG046152. We thank Patricia Purcell, Amy Grayson, and the members of the Kellis lab for their scientific suggestions. Figure 1 was created with the use of <u>BioRender.com</u>. The content is solely the responsibility of the authors.

## Author Contributions

Y.T. conceived, designed, and supervised the study, which was initiated while Y.T. was a postdoctoral researcher in M.K.’s laboratory; W.F.L. performed data analysis and prepared the figures; W.F.L. and N.M. contributed to the biological interpretation of the results with input from Y.T. and M.K.; W.F.L., N.M., and Y.T. drafted the manuscript with input from M.K.; D.A.B. organized reagents. M.K. supervised data access. M.K. and D.A.B. were responsible for funding acquisition, and all authors reviewed and approved the final version of the manuscript.

## Data and Code Availability

We used individual-level data from the Religious Orders Study/Rush Memory and Aging Project (ROSMAP) through the Rush Alzheimer’s Disease Center Resource Sharing Hub (https://www.radc.rush.edu/).

We downloaded the pre-trained polygenic score models characterized from our previous study^68^ from the PGS catalog^122^ (PGS Publication ID: PGP000244) https://www.pgscatalog.org/.

The code used to analyze the data and generate the figures in the manuscript is available at https://github.com/williamfli/AD_PGS_paper_code/tree/main

## Declaration of generative AI and AI-assisted technologies

During the preparation of this work, the authors used ChatGPT and Claude to develop analysis scripts for the project and to refine text. After using this tool/service, the authors reviewed and edited the content as needed and take full responsibility for the results and content of the published article.

