## Supplemental figures and supplemental table legends. for "Dissecting Alzheimer’s disease heterogeneity by cross-trait polygenic prediction"

### Supplemental information

#### Supplemental Figures

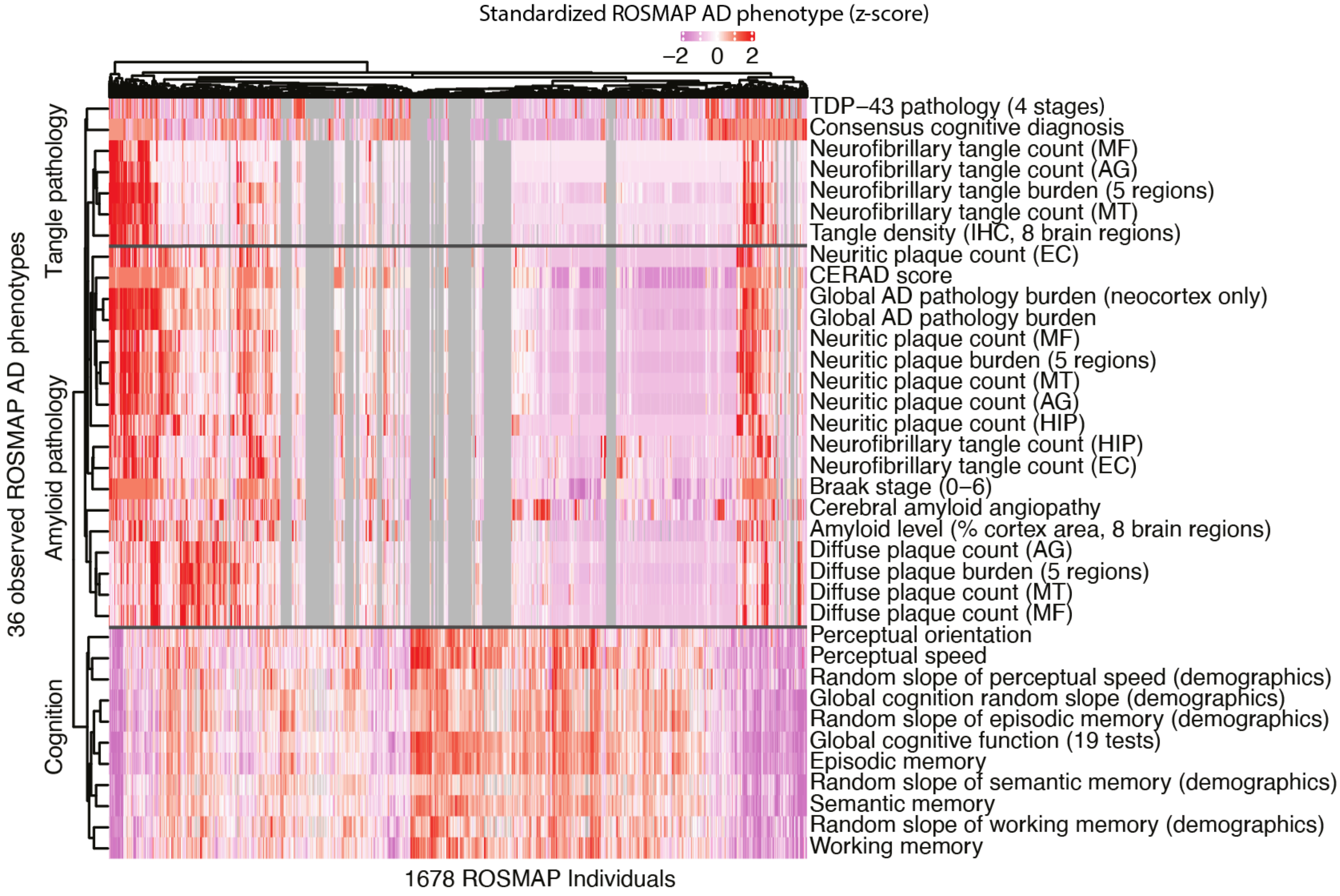


**Figure S1. Phenotypic heterogeneity in the AD phenome in ROSMAP.** We show standardized phenotypes (z-scores) across 1678 ROSMAP individuals (x-axis) and 36 observed ROSMAP AD-related phenotypes (y-axis), each sorted by hierarchical clustering shown as dendrograms. The color represents Z-scores of phenotypic values truncated at 2, with gray indicating missing values. We show three distinct ROSMAP AD phenotype clusters, qualitatively determined to approximately correspond with tangle pathology, amyloid pathology, and cognition.

Abbreviations: TDP-43 pathology (4 stages), transactive response DNA binding protein 43; MF, midfrontal cortex; AG, inferior parietal cortex; MT, midtemporal cortex; IHC, immunohistochemistry; EC, entorhinal cortex; CERAD, Consortium to Establish a Registry for Alzheimer's Disease[42]; HIP, hippocampus; "(demographics)", variable is controlled for demographics.

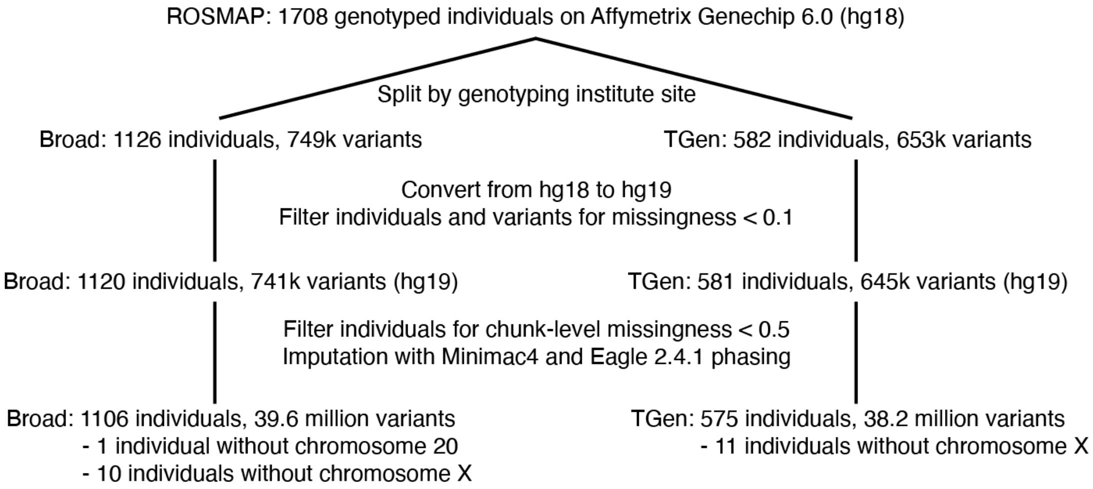


**Figure S2. Quality control of genotype data in ROSMAP.** A flowchart showing the conversion from hg18 to hg19, imputation, and quality control steps on the genotyping data, with the number of individuals and variants at each step. TGen: the Translational Genomics Institute.

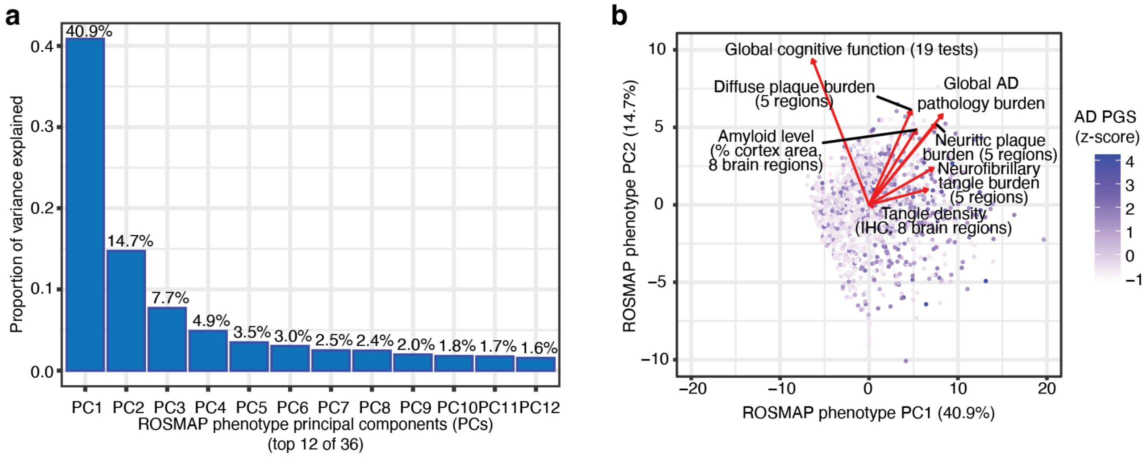


**Figure S3. PCA of ROSMAP phenotypes. (a)** We show variance explained (y-axis) by each of the top 12 ROSMAP phenotype principal components (x-axis) based on a principal component analysis over the individual-level phenotypes. **(b)** We show ROSMAP individuals (scatter) by their principal component (PC) projections onto PC1 (x-axis) and PC2 (y-axis). We color individuals by normalized evaluation of the 36 AD phenotypes across the full dataset, with color scaled by standardized AD PGS (z-scores). We overlay the plot with arrows, with the components of each arrow showing the respective phenotype's contribution to PC1 (x-component) and PC2 (y-component).

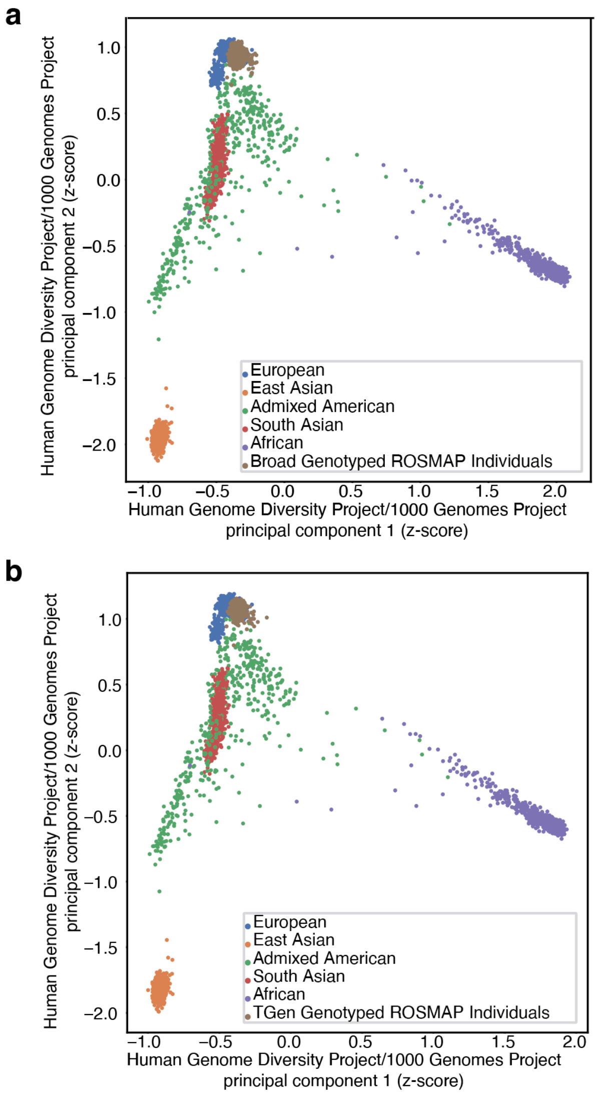


**Figure S4. Quality control of population-level genetic variation.** Individuals genotyped at the Broad **(a)** and the Translational Genome Institute **(b)** are shown in brown and projected onto principal component 1 (x-axis) and principal component 2 (y-axis) from the Human Genome Diversity Project/1000 Genomes Project principal component analysis. Other points represent individuals from the Human Genome Diversity Project/1000 Genomes Project, colored by ancestry. Both sets of genotyped individuals map to the European area of the principal component space.

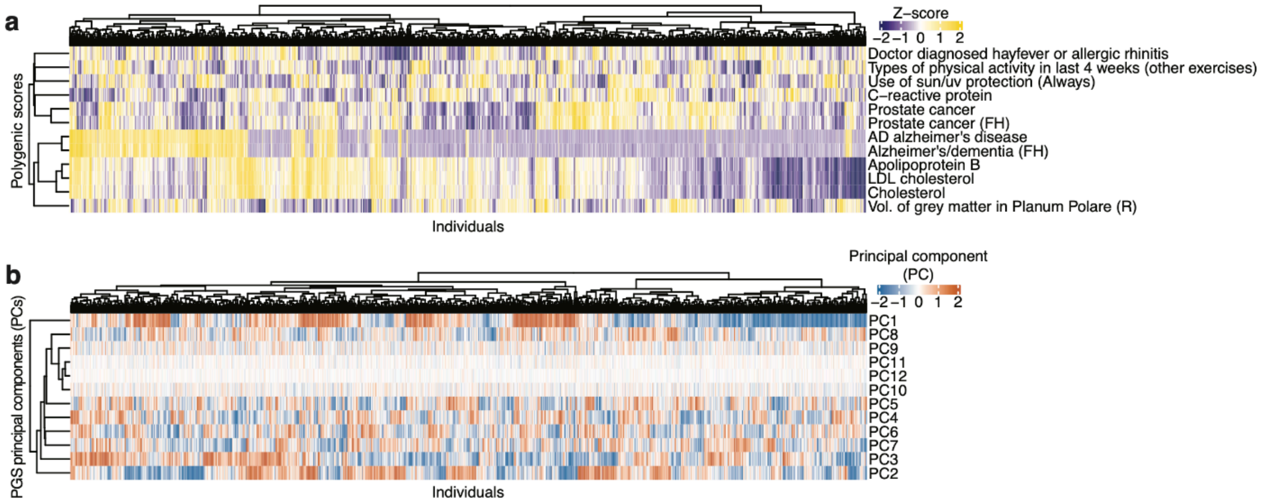


**Figure S5. Individual-level PGS principal component and PGS heatmaps. (a)** Individuals in ROSMAP (x-axis) are shown by their standardized PGS (y-axis, color), with thresholding for an absolute z-score < 2. Dendrograms show hierarchical clustering based on PGS of individuals. **(b)** Individuals in ROSMAP (x-axis) are shown by their polygenic score principal components (y-axis, color), with magnitudes < 2. Dendrograms indicate hierarchical clustering based on individuals' PGS principal component values.

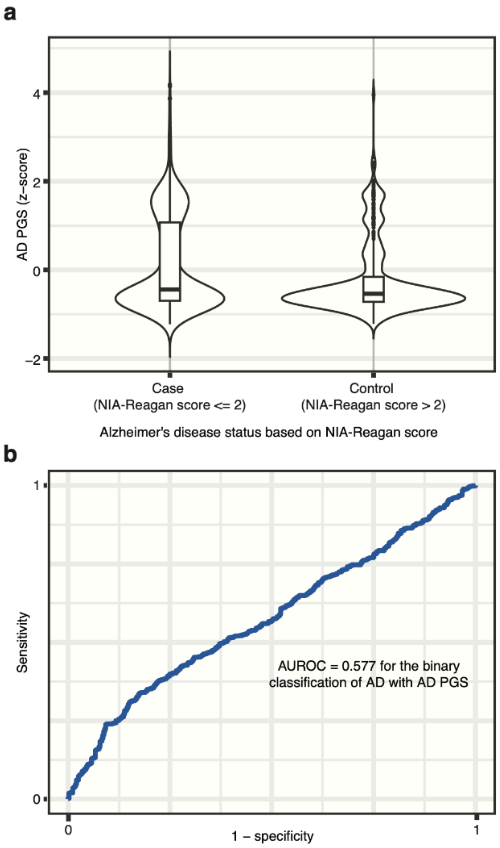


**Figure S6. PGS transferability from the UK Biobank to ROSMAP. (a)** Application of the AD PGS model (y-axis) to predict AD diagnosis (x-axis) based on the NIA Reagan Score diagnostic framework (pathologic diagnosis, performed by a neuropathologist). Box plots show the quartiles of the PGS distributions. **(b)** Receiver operating characteristic (ROC) curve with 1-specificity (x-axis) against sensitivity (y-axis). We report the area under the ROC curve (AUROC).

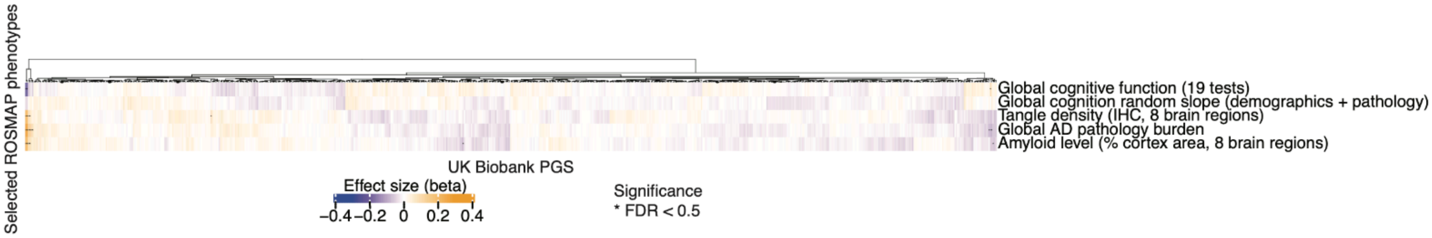


**Figure S7. Stage 1 cross-trait association results for all 713 PGS.** Cross-trait associations between 5 select observed AD phenotypes (y-axis) and all 713 PGS (x-axis). Association sizes are shown by color. All associations with FDR<0.5 are indicated by an asterisk (*).

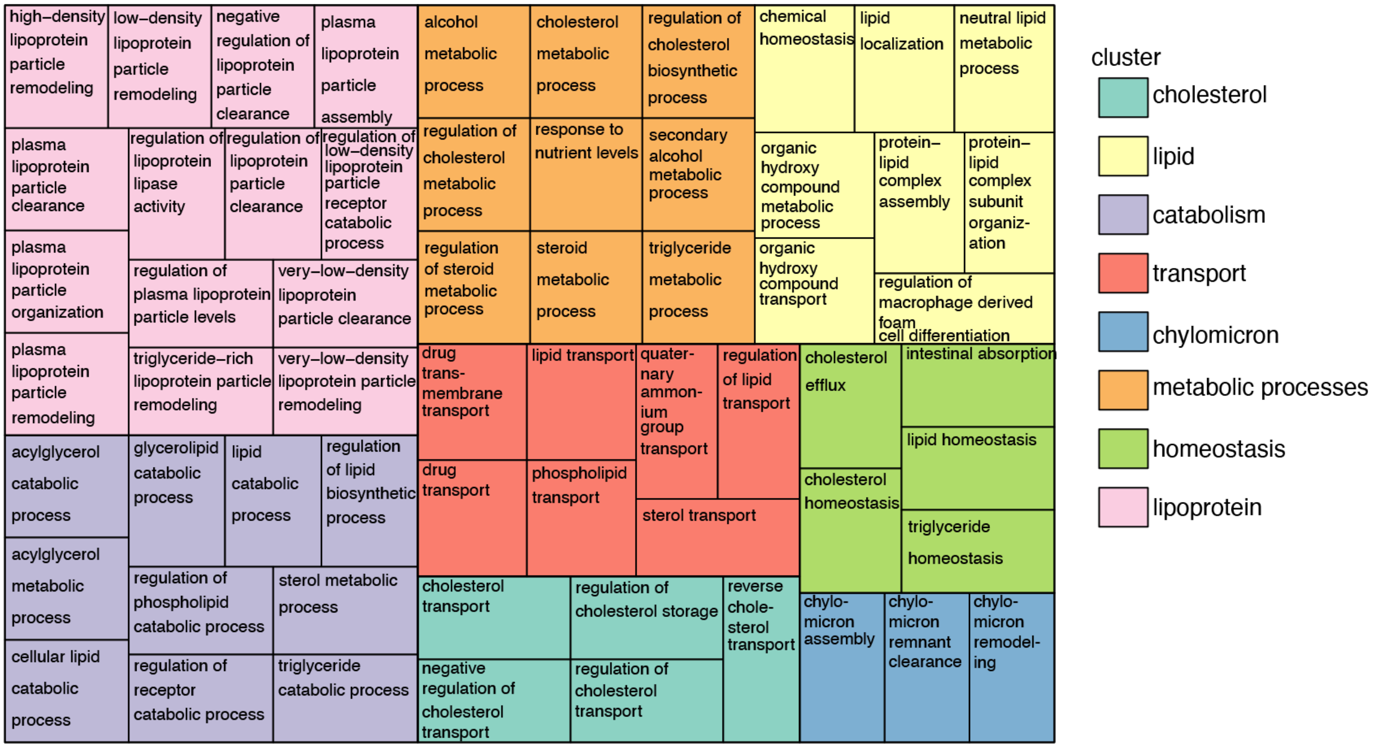


**Figure S8. Semantic organization of Apolipoprotein B PGS model GREAT enrichment Gene Ontology terms.** We present the top 61 Gene Ontology biological process terms for Apolipoprotein B PGS, organized by semantic clustering into 8 biological themes. We selected leading terms to have hypergeometric FDR < 0.05 and binomial FDR < $10^{-15}$.

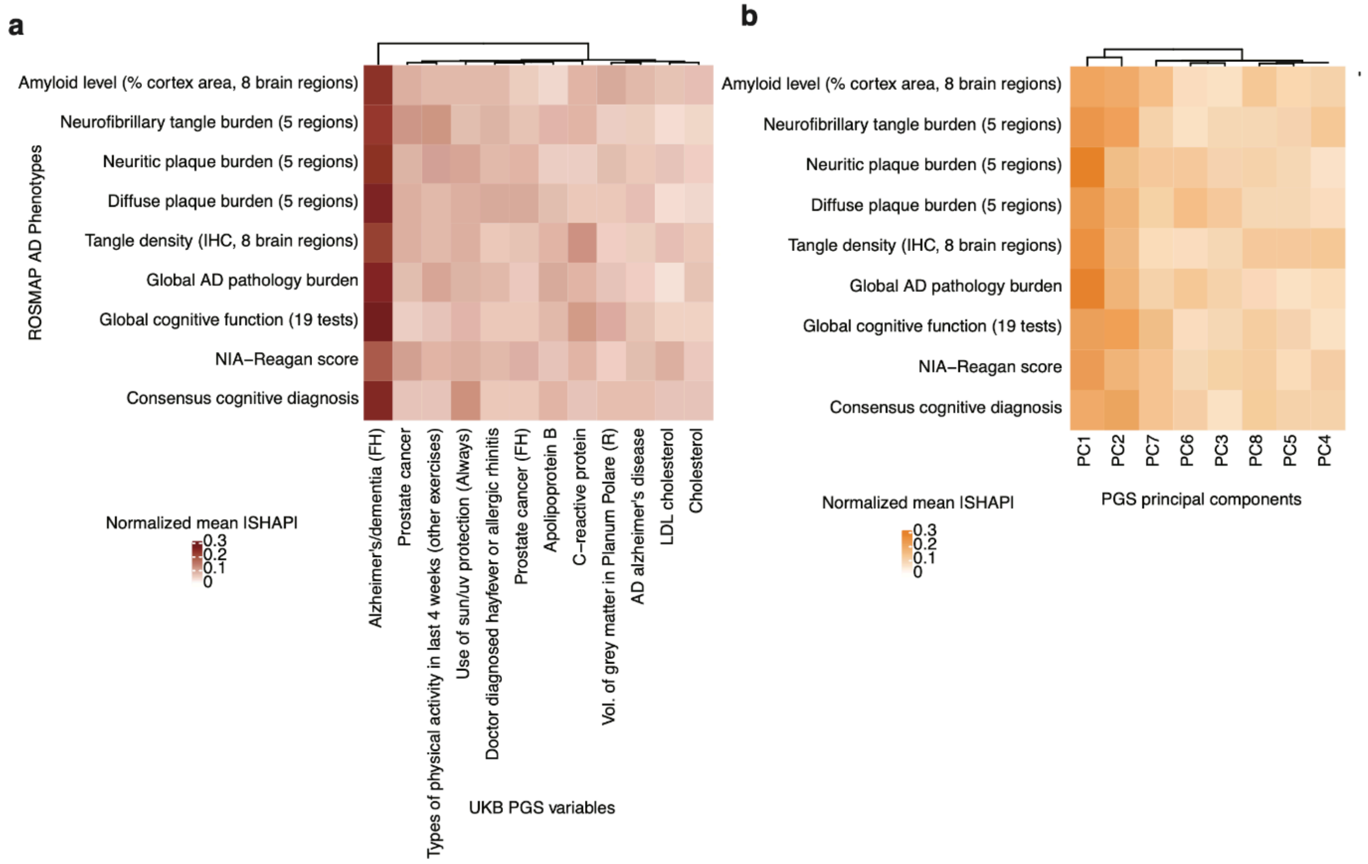


**Figure S9. Predictive contributions (normalized mean absolute Shapley values) of each genetic feature to each phenotype in multiple-PGS genetic prediction models. (a)** Mean absolute Shapley values in the 12-PGS (x-axis) gradient boosted prediction model for each observed ROSMAP AD phenotype (y-axis), normalized so the contribution values for each phenotype (row) sum to 1. **(b)** Mean absolute Shapley values in the 8-principal component (x-axis) gradient boosted prediction model for each ROSMAP AD phenotype (y-axis), normalized so the contribution values for each phenotype (row) sum to 1.

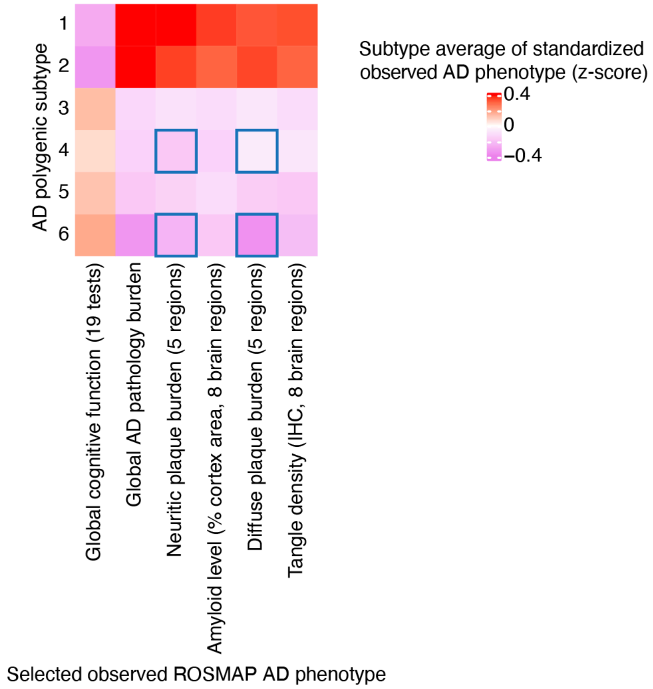


**Figure S10. Heatmap of average standardized AD phenotypes among AD polygenic subtypes.** We show the subtype average (color) of each standardized AD phenotype (column) within each AD polygenic subtype (row). Boxed are the four subtype-specific phenotypes shown in (**Figure 5g**).

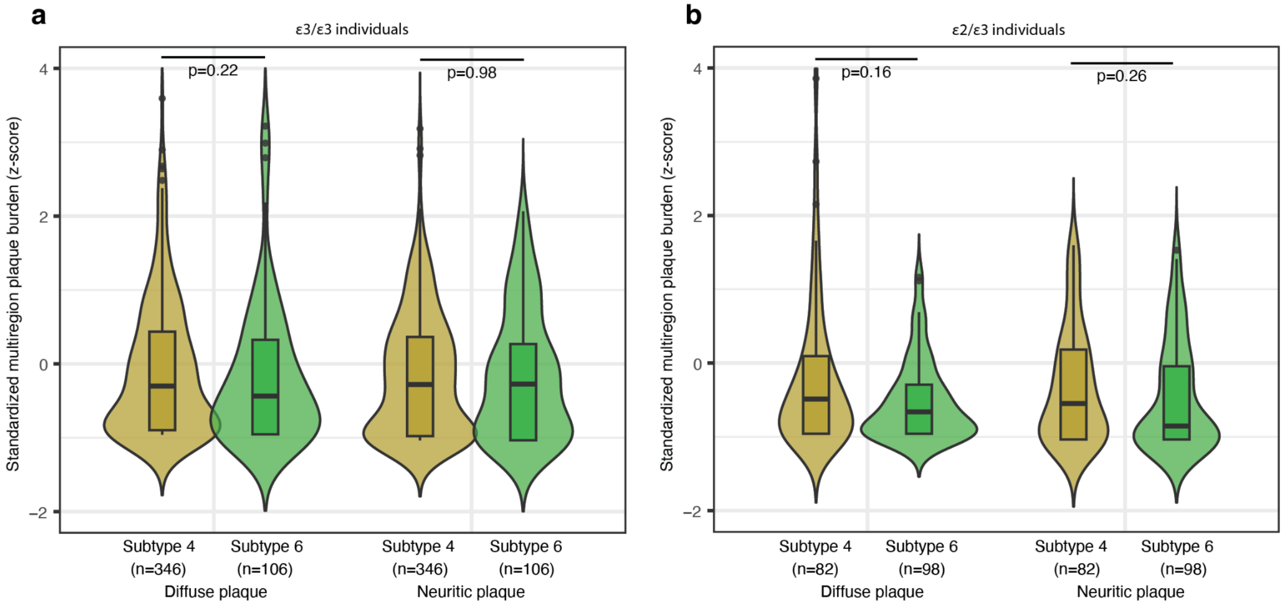


**Figure S11. *APOE*-stratified comparison of observed plaque burdens between AD polygenic subtypes.** We show the distribution of ROSMAP AD phenotype value (color) within each AD polygenic subtype for selected ROSMAP variables (x-axis). We used the Wilcoxon rank-sum test to compare the distributions for each variable (**Methods**). We show this stratified by individuals with ε3/ε3 *APOE* allelotype (**a)** and by individuals with ε2/ε3 *APOE* allelotype (**b**).

#### Supplemental Tables

**Table S1. Observed ROSMAP phenotype variables.** The 126 ROSMAP variables considered in this study. For each variable (row), we indicate whether it is a quantitative phenotype (n=85) and whether it was analyzed as part of the stage 1 analysis (n=5), stage 2 analysis (n=36), or as a covariate (n=3). For each variable (**Observed ROSMAP Phenotype ID**), we show the expanded name (**Observed ROSMAP Phenotype Name**), whether it was used in the stage 1 analysis (**Stage_1**), whether it was used in the stage 2 analysis (**Stage_2**), whether it was used as a covariate (**Covariate**), and whether the variable's data a continuous quantity (**Continuous**).

**Table S2. UK Biobank PGS.** The 713 PGS that were analyzed. Annotated by whether they were analyzed in the Stage 1 analysis or the Stage 2 analysis. For each PGS (**UK Biobank PGS ID**), we show the extended name (**UK Biobank PGS Name**), whether it was used in the Stage 1 analysis (**Stage_1**), and whether it was used in the Stage 2 analysis (**Stage_2**).

**Table S3. Full association results for the Stage 1 analysis (prioritization of AD-relevant PGS using associations with five global observed AD phenotypes).** Each row corresponds to a distinct cross-trait association pair. We show associations between the entire library of 713 UKB PGS and 5 selected ROSMAP phenotypes. For each pair of **UKB PGS** and **Observed ROSMAP Phenotype**, we show the effect size of their association (**Effect Size**), the standard error of the effect size estimate (**Standard Error**), the p-value of the pairwise linear regression (**P-Value**), and the false discovery rate (**FDR**)-corrected q-value over all 3565 (=5 ROSMAP phenotypes * 713 UKB PGS) pairwise associations.

**Table S4. Full association results for the Stage 2 analysis (association of each AD-relevant PGS against all observed AD phenotypes).** Full association results for the stage 2 association. Each row corresponds to a cross-trait pair. We show associations between the 12 prioritized AD-relevant UKB PGS from stage 1 and all 36 AD-relevant ROSMAP phenotypes. For each pair of **UKB PGS** and **Observed ROSMAP Phenotype** we show the effect size of their association (**Effect Size**), the standard error of the effect size estimate (**Standard Error**), the p-value of the pairwise linear regression (**P-Value**), and the false discovery rate (**FDR**)-corrected q-value over all 432 (=36 ROSMAP phenotypes * 12 UKB PGS) pairwise associations.

**Table S5. Association results for the Stage 2 analysis with *APOE* exclusion.** Full association results for the Stage 2 association after excluding the *APOE* region from PGS scoring (**Methods**). Each row corresponds to a cross-trait association pair. We show associations between the 12 prioritized AD-relevant PGS traits from Stage 1 and all 36 observed phenotypes in the AD phenome. For each pair of **UKB PGS** and **Observed ROSMAP Phenotype**, we show the effect size of their association (**Effect Size**), the standard error of the effect size estimate (**Standard Error**), the p-value of the pairwise linear regression (**P-Value**), and the false discovery rate (**FDR**)-corrected q-value over all 432 (=36 ROSMAP phenotypes * 12 UKB PGS) pairwise associations.

**Table S6. Associations of observed ROSMAP AD phenotypes with *APOE*.** Association results of *APOE* against the whole AD phenome of 36 phenotypes. Each row shows the effect size of association, the standard error in the effect size estimate, the p-value, and the false discovery rate. For the genetic feature ***APOE*-gradient** and each **Observed ROSMAP Phenotype**, we show the effect size of their association (**Effect Size**), the standard error of the effect size estimate (**Standard Error**), the p-value of the pairwise linear regression (**P-Value**), and the false discovery rate (**FDR**)-corrected q-value over all 36 associations.

**Table S7. Apolipoprotein B PGS model GREAT enrichment.** Results from GREAT enrichment analysis of the top 1000 weights for the PGS model for Apolipoprotein B. Each row indicates enrichment statistics for a GO biological process. For each **Gene Ontology Term**, we show the binomial FDR-corrected q-value of the GREAT enrichment (**BinomFdrQ**), the region-fold enrichment (**RegionFoldEnrich**), and the hypergeometric FDR-corrected q-value of the enrichment (**HyperFdrQ**).

**Table S8. Prostate cancer PGS model GREAT enrichment.** Results from GREAT enrichment analysis of the top 1000 weights for the PGS model for prostate cancer risk. Each row indicates enrichment statistics for a GO biological process. For each **Gene Ontology Term**, we show the binomial FDR-corrected q-value of the GREAT enrichment (**BinomFdrQ**), the region-fold enrichment (**RegionFoldEnrich**), and the hypergeometric FDR-corrected q-value of the enrichment (**HyperFdrQ**).

**Table S9. AD phenotype prediction by different genetic features.** The expanded table of Pearson correlation statistics on the test set for XGBoost-based predictive models for observed AD phenotypes trained on different genetic features (see Table 1). Each row corresponds to a different predicted **Global Observed ROSMAP Phenotype**, and each column to a different genetic feature or set of genetic features. Each cell is the Pearson correlation coefficient on the test set for that genetic feature-phenotype pair.
